# Aggregates dramatically alter fibrin ultrastructure

**DOI:** 10.1101/432138

**Authors:** X. García, L. Seyve, Z. Tellier, G. Chevreux, N. Bihoreau, B. Polack, F. Caton

## Abstract

Among the many factors influencing fibrin formation and structure (concentration, temperature, composition, pH,…), it has been suggested that the polydispersity of fibrinogen may play an important role. We propose here a detailed investigation of the influence of this parameter on fibrin multiscale structure.

Two commercial fibrinogen preparations were used, a monodisperse and a polydisperse one. First, the respective compositions of both fibrinogen preparations were thoroughly determined by measuring the FXIII, fibronectin, α,β and γ intact-chains contents, theγ/γ’ chains ratio, the N-glycosylation and the post-translational modifications. Slight variations between the composition of the two fibrinogen preparations were found which are much smaller than the compositional variations necessary to alter significantly fibrin multiscale structure as observed in the literature. Conversely, MALLS coupled SEC and DLS measurements showed that the polydisperse preparation contains significant amounts of aggregates while the other preparation is essentially monodisperse.

The multiscale structure of the fibrins produced from those two fibrinogen preparations was determined by using X-ray scattering, spectrophotometry, and confocal microscopy. Results show that fibers from the monodisperse fibrinogen present a crystalline longitudinal and lateral structure and form a needle-like network. The internal structure of fibers produced from the polydisperse fibrinogen looks amorphous with star-like branching nodes. The multiscale structure of mixtures between the two preparations shows a smooth evolution, demonstrating that the quantity of aggregates is a major determining factor for fibrin multiscale structure. Indeed, the effect of a few percent in mass of aggregates is larger than any other effect due to compositional differences under the same reaction conditions. Finally we propose a mechanistic interpretation of our results which points at a direct role of the aggregates during polymerization which disrupt the ideal ordering of monomers inside fibrin protofibrils and fibers.

## 1 Introduction

The formation of the fibrin clot is essential in the process of blood coagulation. Fibrin forms a protein scaffold that enables the organism to close off damaged blood vessels. In the first step of fibrin formation, the fibrinopeptides A and B from the protein fibrinogen are cleaved by thrombin, producing the so-called fibrin monomers which then polymerize into a fibrin network. Fibrinogen itself is a plasma 340-kDa centrosymetric protein. Its structure is constituted by three aligned domains. The central part of the molecule (E-region) is slightly smaller (c.a. 5nm of diameter) while the distal regions (D-regions) present a 6nm diameter to form a structure about 45nm in length.

While some aspects of the formation of the fibrin network from fibrin monomers are still under debate, it is known that various genetic and environmental factors influence not only the fibrin structure and function but that they can also be related with thrombotic disease (e.g. 1). Indeed, many clinical studies have associated the fibrin properties with thrombosis (2, 3). Among the many factors influencing fibrin polymerization, structure and function (ionic strength, pH, concentrations, γ’ content … (4-9)), Huang et al. (10,11) suggested that the size dispersity, i.e. the amount of aggregates or oligomers present in the fibrinogen preparation, may play an important role. Indeed, fibrins made from chromatography fractionated fibrinogen exhibited final turbidities between two and three times higher than the unfractionated samples (10), and significant differences both in kinetics as well as in the apparent size of the polymerizing objects were observed in a later study (12). However, a Small Angle X-Ray Scattering study of this fractionation process showed significant in–column degradation (13) and gel filtration could change the concentration in important proteins present in the fibrinogen preparations (called co-purified proteins), such as fibronectin, FXIII, … Likewise, fibrinogen isoforms (glycoforms, γ’ content, oxidized forms, N-/C-terminus cleaved forms, …) are likely to be prone to variability during the chromatography fractionation process and therefore to behave specifically *vis-à-vis* polymerization.

As the relative compositions of the mono- and poly-disperse fibrinogen preparations were not characterized nor the molar masses of the different fractions measured, the question of whether the observed effect is a consequence of variations in aggregate content and/or composition remains open. Furthermore, single-wavelength optical-density measurements only indicate an overall change in fibrin structure but provide no indication about the morphological changes or the scale(s) affected, nor about the involved mechanisms.

In the last decade or so, about 75% of the fundamental work on fibrin formation has been performed on fibrinogen from Enzyme Research Laboratories (see Supporting Information (SI) S1), hence with rather constant compositional and polydispersity profiles. While this is pertinent for comparison purposes, the effect of fibrinogen’s polydispersity may have been entirely overlooked in the current fibrin-related literature, while it is a well-known factor in standard polymer science (e.g. 14).

The above observations raise four important questions. First, does fibrinogen dispersity really influence the multiscale structure of fibrin, or were Huang et al. observations (11) a consequence of the purification procedure they used? Second, what are the scales affected by dispersity and how are those scales affected? Third, are those differences also observed at physiological fibrinogen concentrations (2-4 mg/ml) or limited to the very low concentrations used in previous studies? Finally, are the dispersity-induced structural effects important? In other words, are they smaller or larger than those induced by varying compositional aspects of the fibrinogen preparations?

In the following, we start by describing the numerous experimental methods used in this work. To characterize the two fibrinogens, we performed Refractive index and MALLS-coupled Size Exclusion Chromatography, Dynamic Light Scattering, Reverse phase chromatography coupled to mass spectrometry, and all necessary co-purified proteins quantification assays. The methods used to determine the structure of the fibrins at each scale are then described, starting with Small Angle X-Ray Scattering, Spectrophotometry, and finally Confocal Microscopy.

Then, we present the detailed characterization of the physical and physicochemical properties of the different fibrinogens, showing that the two fibrinogens used in this study present identical compositional profiles while differing strongly in their aggregate content. We then investigate the effect of this size dispersity on the nano- and micro-scale structure of the fibrin, showing that the two fibrins present vastly different structures, at all scales. We finally discuss the present findings in the light of recent results concerning the effect of compositional and isoform changes on fibrin structure and propose a mechanistic explanation of our results.

## 2 Materials and Methods

### Materials

Human thrombin was purchased from Cryopep (Montpellier, France) as a 12 μL solution containing 298,9 IU. The solution was diluted to 200 IU/ml in a 2-(N-morpholino)ethanesulfonic (MES) buffer (20mM MES, 50mM NaCl, pH 6.5) aliquoted and kept frozen at −80°C.

Two fibrinogens were used: Clottafact (Laboratoire Français du Fractionnement et des Biotechnologies, Les Ulis, France) and Fib1 (Enzyme Research Laboratories, Swansea, UK). Fibrinogens were reconstituted using manufacturers’ guidelines: Fib1 and Clottafact fibrinogens were reconstituted by adding respectively 25 mL and 100 mL of Sterile Water for Injection into the product vials. Then, the product vials were incubated at 37° C and gently swirled until the product was fully dissolved. Both fibrinogens were dialyzed together twice overnight against >100 volumes of HEPES buffer (140 mM NaCl, 20 mM HEPES, 5 mM CaCl_2_, pH 7.4), aliquoted and kept frozen at −80° C. Fibrinogen concentrations were determined by absorbance at 280 nm using a specific absorption coefficient of E^280^= 1.51 ml⋅mg^-1^⋅cm^-1^.

Dithiothreitol (DTT), iodoacetamide (IAM), urea, phosphate buffered saline (PBS), ammonium carbonate, and HEPES were purchased from Sigma-Aldrich Chemical (St. Louis, MO, USA). Acetonitrile (MeCN) was HPLC reagent grade and purchased from JT Baker (Philipsburg, NJ, USA). Trifluoroacetic acid (TFA) was from Merck Biosciences (Darmstadt, Germany). All the aqueous solutions were prepared using ultra-pure water (18.2 MΩ-cm resistivity at 25°C, total organic carbon (TOC) < 5 ppb).

### Sample preparation

Fibrinogens were thawed at 37°C for 5 minutes and equilibrated at room temperature for another 5 minutes before use, filtered with a 0.2 μm ClearLine syringe filter. Concentrations were then determined by absorbance at 280 nm, and adjusted as desired. Thrombin was thawed at 37°C for 1 minute, diluted in HEPES buffer and immediately used.

Fibrin clots were formed by incubating fibrinogen (typically 0.5, 1 and 3 mg/ml) with 0.1 IU/ml thrombin (final concentration) at 37°C during 90 minutes.

### Size exclusion chromatography

The molecular distribution of the different fibrinogens was analyzed using a Superose 6 column (GE Healthcare, USA) on an Elite LaChrom system (Hitachi, Japan), coupled with a Dawn Heleos II - Multi-Angle static Light Scattering system (Wyatt Technology, Santa Barbara, USA) and a Optilab T-rex – Refractometer (Wyatt Technology). The same HEPES buffer as before was used. Fifty microliters of fibrinogen (5 mg/ml) were applied and the column was developed at a 0.5 ml/min flow rate. Elution (λ=280 nm), MALLS and Refractometer profiles were analyzed using the ASTRA software from Wyatt Technology.

### Dynamic Light Scattering

Dynamic Light Scattering measurements were performed using a CGS-8FS/N069 apparatus from ALV Technology (Manila, Philippines) with a 35 mW, 632.8nm laser from JDSU (Milpitas, CA, USA). Fibrinogens (1 mg/ml) were loaded in 10 mm diameter cylindrical cells, immerged in a toluene bath at 25.0 ± 0.1 °C. Data were collected at 90° for 120 s. Hydrodynamic radii distributions were determined using Contin analysis.

### Protein assays

Fibronectin level was assayed on a Siemens BNII nephelometer. Briefly, the sample is mixed with a polyclonal rabbit anti-Human fibronectin to generate immune complexes measured by nephelometry. The fibronectin concentration is deduced by interpolation with a standard curve using the Dade Behring N protein standard PY.

FXIII levels were determined by ELISA using sheep polyclonal anti-Human FXIII (Cedarlane, CL20057A) for the coating and the same polyclonal anti-Human FXIII conjugated with horseradish peroxidase (Cederlane, CL20057HP) as secondary antibodies. Standard human plasma (Siemens, ORKL 17) was used to establish the calibration curve. Results are expressed as International Unit (IU) knowing that 1IU is equivalent to 30 μg/mL.

### Reverse phase chromatography coupled to mass spectrometry (MS)

Fibrinogen (100 μg) was vacuum-dried and dissolved in 35 μL of a 8 M urea and 0.4 mM ammonium carbonate solution at pH 8.0. Reduction was done by adding 10 μL of a 40 mM DTT solution in water and incubating the resulting mixture for 20 min at 50°C. After cooling at room temperature, 10 μL of a 80 mM IAM solution in water were added and the solution was incubated at room temperature for 20 min in the dark. Reverse phase high-pressure liquid chromatography (RP-HPLC) was performed using an ACQUITY UPLC system (Waters, Milford, MA, USA). An amount of 20 μg of sample was injected on a Pursuit 3 diphenyl reverse phase column (150×2.0 mm, 3 μm, Agilent, Santa Clara, CA, USA) equilibrated at 70°C and operated at a flow rate of 200 μL/min. An aqueous solution containing 0.1% TFA and MeCN containing 0.1 % TFA were respectively used as buffer A and buffer B; proteins were eluted by using an increasing gradient of buffer B. After separation, reduced and alkylated fibrinogen chains were detected by UV at 280 nm and MS analysis was achieved by interfacing the UV detector output to a SYNAPT G2-S HDMS mass spectrometer (Waters, Milford, MA, USA) scanning from *m/z* 500 to 2000.

### Small angle X-ray Scattering

Small Angle X-ray Scattering (SAXS) experiments were performed at the ID02 line at the European Synchrotron Radiation Facility (ESRF, Grenoble, France). The samples (2 volumes of fibrinogen and 1 of thrombin) were mixed directly in a 12mm x 12mm x 3mm home-made cell with 20 μm mica windows, thermostated at 37± 0.3°C (16). The sample-to-detector distance was set to 7 m and the acquisition time was set to 0.1 s. To avoid radiation-induced degradation of the protein, the cell was displaced by a motorized x-y stage between each measurement in a snake-like fashion, by 2 mm horizontally and 2 mm vertically at the end of a line, i.e. each time by a distance much larger than the beam size (0.5×0.5 mm). The results presented here are the average of the 15 last acquisition times corresponding to steady state scattering curves. The constancy of the signal over several centimeters of the cell shows that the polymerization was finished and that the initial mixing was good. A comparison between two replicate experiments is presented in SI S5, fig. SI.6.

### Spectrophotometry

Fibrin gels were formed in 96-well Immulon 2HB plates (Fischer scientific, Illkirch, France) by mixing 2 volumes of fibrinogen (120 μl) and 1 of thrombin (60 μl). Optical density spectra (500 nm to 800 nm) were measured after 90 minutes at 37 ± 0.3 °C using a SpectroStar Omega (BMG Labtech Ortenberg, Germany). For the total volume used in the experiments, the path length was 4.38 mm as determined from calibration using a range of known absorbances.

Optical density data were analyzed using a corrected version of Yeromonahos’s (7,15,16) model. See SI S4 for a discussion concerning the model choice and pertinence. Each data point represents the mean and standard deviation calculated from at least 12 individual experiments.

### Confocal Microscopy

Microscopic images of fibrin networks were obtained using Alexa488 fluorescent fibrinogen (Invitrogen, Breda, Netherlands) mixed with the unlabelled fibrinogens at a 1:10 ratio. Each mixture was filtered with a 0.2 μm ClearLine syringe filter and the concentrations were then determined by absorbance at 280 nm. Samples were polymerized using 0.1 IU/ml of thrombin and various fibrinogen concentrations (0.5, 1 and 3 mg/ml). Fibrinogen and thrombin were mixed in Eppendorf Tubes 3810X, then rapidly injected in Secure Seal Hybridization Chambers (Grace Biolabs, Bend, OR, USA) and polymerized during 60 minutes at 37° C. Confocal image stacks of the fibrin networks were acquired using a Zeiss LSM710 confocal microscope with a 63x/1.2 water immersion objective (Zeiss, Oberkochen, Germany). The 3D stacks were 67.5 μm in the x-y direction and 25 μm in the z direction with resolutions of 100nm and 250nm respectively. The confocal image stacks of the fibrin networks are homogeneous both in the imaging plane and also in depth (data not shown).

For the visualization and presentation of those stacks, we used a recently developed ImageJ plugin: Smooth Manifold Extraction (SME) (17). This method, unlike z-maximum intensity projections, provides a robust 2D representation of 3D objects which preserves local spatial relationship, in particular branching (see SI S6.1 for details and examples). Finally, the pore size was determined using the bubble method (18, 19). For details about the segmentation and pore size analysis, see SI S6.2.

## 3 Results

### 3.1 Physico-chemical characterization

#### Molecular size distribution

The SEC elution profiles of the two fibrinogens are presented in Fig. 1. They both show a main peak at an elution volume of 11ml, as well as a small broad peak around 9.3 ml, albeit with different intensities (Fib1 > Clottafact). In addition, Fib1 presents a well-defined high molecular weight peak occurring at an elution volume of about 7.5 ml. The analysis of the MALLS and index refraction profiles obtained simultaneously to the elution profile (see material and methods) yielded the molar masses of the species corresponding to each of those elution peaks as well as their relative concentrations.

**Figure 1.**
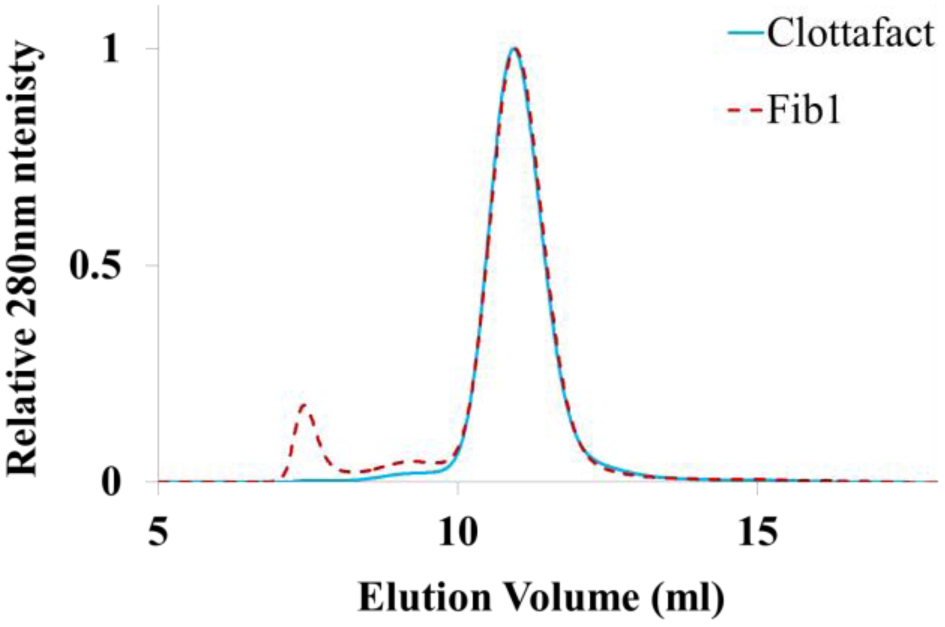
Molecular size distribution of Clottafact and Fib1 determined by size exclusion chromatography.

A (weight-averaged) molar mass of 325 kDa is obtained for the 11 ml elution peak observed in both fibrinogens, which corresponds closely to the molar mass of fibrinogen monomers (e.g. 13). The ratio of the weight-averaged mass to the number-averaged mass (Mw/Mn) for this peak is below 1.01, indicating good monodispersity. For the broad small peak at 9.5 ml also observed in both fibrinogens, a molar mass of about 1 MDa is obtained, with a significant polydispersity, corresponding probably to a distribution of oligomers. Finally, a mass of about 5 MDa can be assigned to the large peak observed at 7.5 ml in Fib1, corresponding most likely to fibrinogen aggregates. The elution profile and molar masses obtained for Fib1 are in excellent agreement with those of Brookes et al. (13, fig. 3) which were also obtained on Fib1 fibrinogen, but using a size exclusion chromatography column coupled to a small angle X-ray scattering cell.

Those results demonstrate the presence of both oligomers and large aggregates in Fib1 while the Clottafact preparation is close-to-monodisperse with a small amount of oligomers and no detectable aggregates. The relative mass of the aggregates present in Fib1 is found to be of about 4% of the total mass of fibrinogen.

#### Dispersity analysis by Dynamic Light Scattering

Dynamic Light Scattering (DLS) was used to confirm the above results by determining the average hydrodynamic radius of the different fibrinogens. Since the SEC-MALLS results showed a tri-disperse size distribution, we tried to use a triple exponential fit of the raw correlation spectra to obtain the hydrodynamic size distribution of the molecules. However, those fit were not robust as the result was strongly dependent on the initial values used for the fit. Therefore, we only present the average hydrodynamic radii and the polydispersity indexes obtained by a Contin analysis.

Clottafact fibrinogen has a hydrodynamic radius of 11nm, a hydrodynamic size corresponding well to that of fibrinogen monomers (11,12,20,21). The measured polydispersity index is of 25%, close to the value of 20% below which samples are considered truly monodisperse. Conversely, Fib1 fibrinogen has a hydrodynamic radius of 22nm, with a polydispersity index of 52%. This result confirms that Fib1 fibrinogen presents mixtures of higher molecular weight species.

#### RP-HPLC-Mass Spectrometry analysis

The two fibrinogens were further analyzed by RP-HPLC-Mass Spectrometry after reduction and separation of the α, β, and γ chains. MS was used to determine the sequence integrity of the different chains as well as their post-translational modifications. The resulting chromatograms (UV detection) for the two fibrinogen preparations are essentially identical in all respects (see SI2.2 and fig. SI.2 for a description and summary of the MS based identifications).

Finally, SDS-PAGE of reduced fibrinogens showed similar γ’/γ chains ratio with 14% for Clottafact and 10% for Fib1 (data not shown).

#### Co-purified proteins content

Co-purified proteins (FXIII and fibronectin) concentrations measured as described in materials and methods are shown in table 1 (see also SI2.1 and fig. SI.1). The main finding is that Clottafact and Fib1 have almost identical composition in co-purified proteins. The only slight difference is the fibronectin content (0.05 vs 0.02 mg/mg) which is low compared to the normal plasmatic value (0.12 mg/mg), and much smaller than the range for which significant structural variations are observed in the literature (0.8 mg/ml) (21, 22) (see SI3, Table SI3.1).

**Table 1:** Physico-chemical characteristics of the two fibrinogens.

|  | Clottafact | Fib1 |
| --- | --- | --- |
| Presence of aggregates | Not detectable | 4% in total Fg mass |
| Average hydrodynamic radius (nm) | 11 | 22 |
| FXIII (U/mg Fg) | 0.26 | 0.25 |
| Fibronectin (mg/mg Fg) | 0.05 | 0.02 |
| Post-translational modifications | identical | Identical |
| Intact α-chains (% vs Clottafact) | 100% | 95% |
| Respective intact $\beta$ and $\gamma$ chains | 100% | 100% |
| $\gamma'/\gamma$ chains ratio (%) | 14 | 10 |

The results of the physico-chemical analyses are summarized in table 1 and show that the composition of the two fibrinogens is almost identical except for the presence of large molecular weight aggregates.

### 3.2 Fibrin’s ultrastructure

Since polydispersity could influence fibrin structure at the molecular, fiber and network scales, each scale was investigated with the appropriate tool (see figure 2).

**Figure 2:**
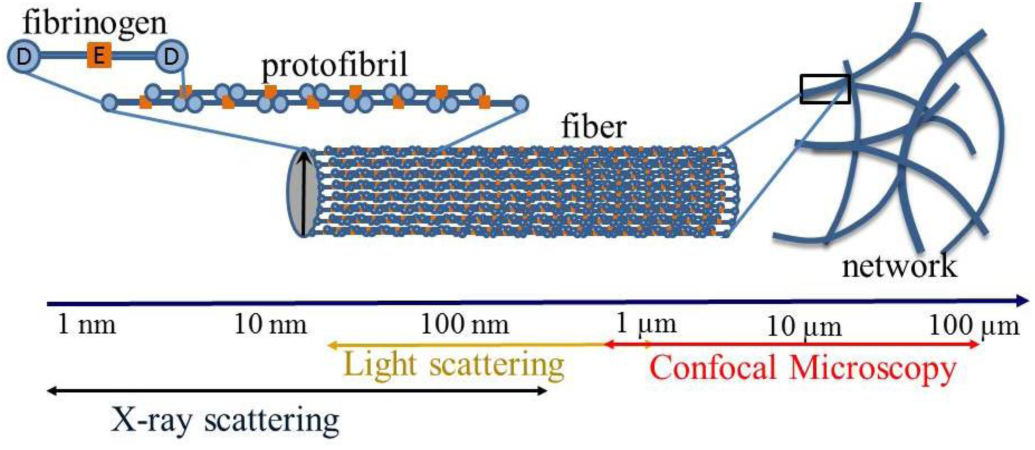
Multiscale structure of the fibrin clot.

#### Fibrin fibers internal structure

We used Small Angle X-Ray Scattering to determine the molecular organization inside fibrin fibers. Figure 3A shows X-ray intensity spectra obtained on mature fibrin clots. The general shape of the scattering curves is identical to those available in the literature (7,18). A replicate experiment was performed, yielding an identical spectrum (see SI5 and fig. SI.6), showing the excellent reproducibility of the protocols and methods. Spectra from each fibrin gel showed a main peak at a *q-*vector ~ 0.3 nm^-1^ corresponding to the usual 22.5 nm periodicity of half-staggered fibrin monomers. However, significant differences are observed around this peak, *i.e.* between 0.2 and 0.5 nm^-1^

**Figure 3.**
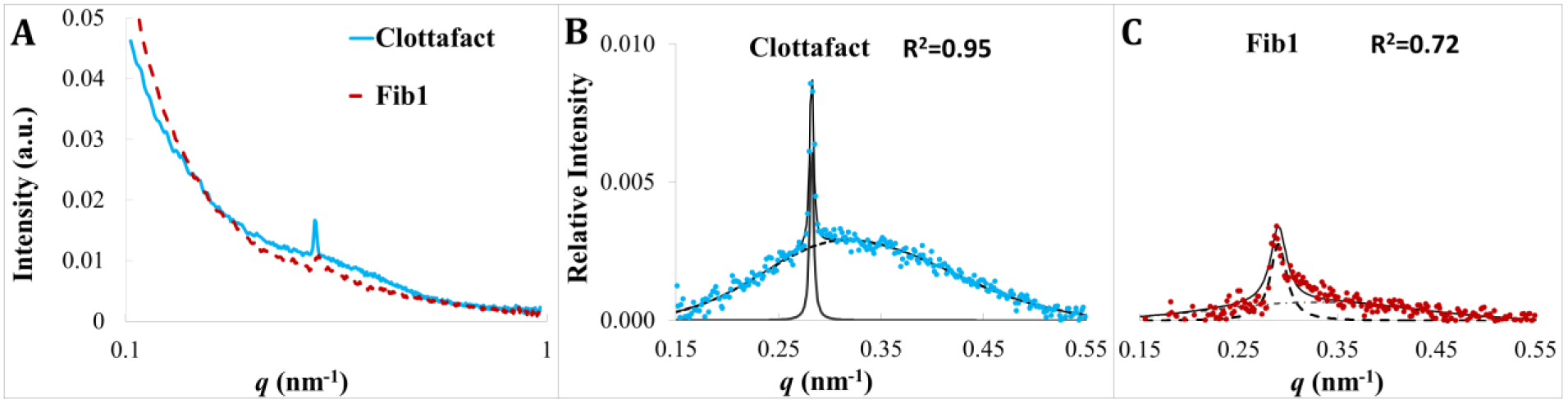
Relative intensity of small angle X-ray scattering for fibrin clots around the 22.5nm periodicity peak. SAXS was performed as described in material and methods. The relative intensity was determined according to the method of Missori et al. (23). Clots were formed using 1 mg/ml of each of the two fibrinogens coagulated using 0.1 IU/ml of thrombin. For B and C, continuous lines are the total fit, while dashed lines are the individual peaks. The R2 values are the goodness-of-fit coefficients.

To isolate those differences, we adapted Missori *et al*. (23) method to our data: the log-log scattering curves were first fitted with an order 3 polynomial. This polynomial was then subtracted to the scattering data, yielding the curves shown in figure 3B and 3C. The data were fitted to two Gaussian distributions (23), one for the main ~22.5nm peak and the other Gaussian for the broad peak (dashed lines).

It is obvious both in figure 3B and in figure 3C that Clottafact fibrin exhibits a much higher amplitude and narrower main peak than Fib1 fibrin. Clottafact fibrin also shows a well-defined secondary broad peak that is not observed for Fib1 fibrin (Fig. 3C).

Clottafact (Fig. 3B) sharp peak is at *q_1_* = 0.284 nm^-1^ with a full-width-at-half-maximum (FWHM) of 0.0026 nm^-1^ and the second broader peak is at *q_2_* = 0.327 nm^-1^ with an FWHM of 0.094 nm^-1^. The sharp peak can be assigned to a Bragg diffraction from a repeat distance of d_1_=2π/q_1_=22.2 nm that corresponds well to the usual periodicity of the half-staggered arrangement of the protofibrils (see fig. 2). Its FWHM gives a persistence length of 2.4 μm for this longitudinal feature (i.e. along the length of the fiber). This indicates that the fibers are straight over this length and without defects in the 22.2 nm half-staggered arrangement. The second, broader peak can be assigned to a repeat distance of d_2_=2π/q_2_=19.1 nm which correspond accurately to the lateral dimension of the unit cell of the protofibrils crystal described theoretically by Yang *et al.* (24) and very close to the one observed experimentally by Freyssinet *et al.* (25). This result suggests that the fibers are well organized laterally (see fig. 2).

Fib1 fibrin (Fig. 3C) shows a small main peak at about 0.285 nm^-1^ *i.e.* a longitudinal repeat of 22.1 nm with a FWHM of ~0.01 nm^-1^ giving a persistence length of ~0.63 μm, about four times smaller than for Clottafact. Thus, fibrin fibers made from Fib1 show a 22.5 nm repeat that is either much less regular (less crystalline) and/or that these fibers are much shorter or much curvier than Clottafact’s. The lateral structure of those fibers is too weak to be analyzed, suggesting also a much less regular internal lateral structure. Those results indicate that the molecular organization of fibrin fibers made from Clottafact is close to crystalline both longitudinally and laterally while the fibrin fibers made from Fib1 are much less organized.

To extend the scope of the investigation, we performed spectrophotometry measurements (7) for a wider range of experimental conditions and numerous replicates, experiments that would have been unacceptably time-consuming on a synchrotron. Figure 4A shows that fibers from Clottafact clots have about twice as many protofibrils than those from Fib1. This result holds from 0.4 to 3 mg/ml, i.e. up to the normal physiological range. The spectrophotometry method also determines the average radius of the fibers from which the protein density within the fibers can be deduced from ϱ=μ/(πr^2^) (e.g. 7). Figure 4B shows that the protein density inside Clottafact fibrin fibers is significantly larger (30-50%) than the density of fibers from Fib1. The range of density values obtained here are in agreement with previous work (15, 26). The validity of the spectrophotometric model was further verified by confronting the internal density of the fibers to confocal pore size measurements (see SI S7 and fig. S10 for details).

**Figure 4.**
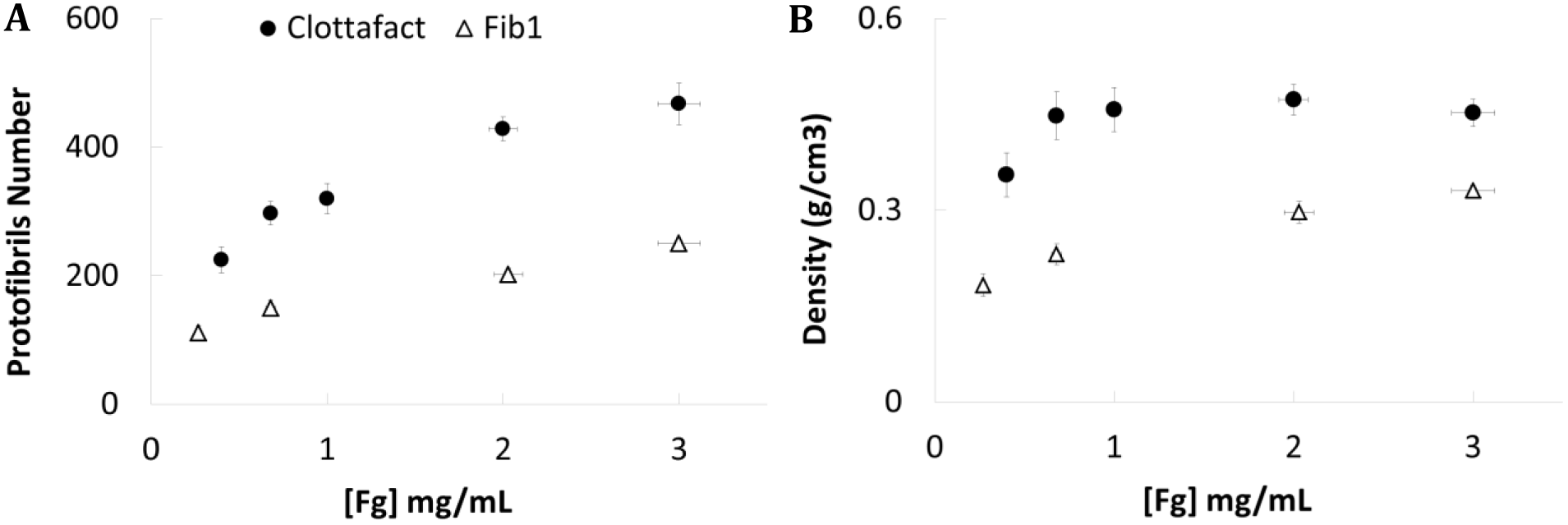
Number of protofibrils and protein density inside fibrin fibers. **A.** Final number of protofibrils vs fibrinogen concentration **B.** Protein density in the fibers vs fibrinogen concentration. Clots were formed using several concentrations of each of the two fibrinogens and coagulated using 0.1 IU/ml of thrombin.

To summarize the nanostructural findings, SAXS indicates that Clottafact fibrin fibers are close to crystalline whereas the internal structure of fibers produced from Fib1 fibrinogens is considerably less organized, *e.g.* amorphous or fractal as was advocated previously (7, 27, 30). Spectrophotometry results point in the same direction showing that Clottafact fibers possess a significantly higher internal density than Fib1 fibers.

#### Fibrin microstructure: Confocal microscopy

Finally, to check whether the fibrinogen preparation differences could also impact the microstructure, we investigated the network organization of fibrin clots at the micron scale. The 3D microstructure of the different fibrins is visualized using the recently developed “Smooth Manifold Extraction” (17). The main asset of this method is that, unlike maximum intensity projections, it does not create spurious intersections (see SI6.1 for a comparison between the two methods). Major differences in the fibers geometry are observed in fig.5: Clottafact fibrin shows needle-like fibers that are straight over several μm, while Fib1 produces networks with much shorter and curved fibers. The observed straightness of Clottafact fibers is in perfect agreement with the very large persistence length obtained in the SAXS measurements while Fib1 fibers shortness or snake-like characteristic is also in agreement with the much lower persistence length obtained from the SAXS measurements.

**Figure 5.**
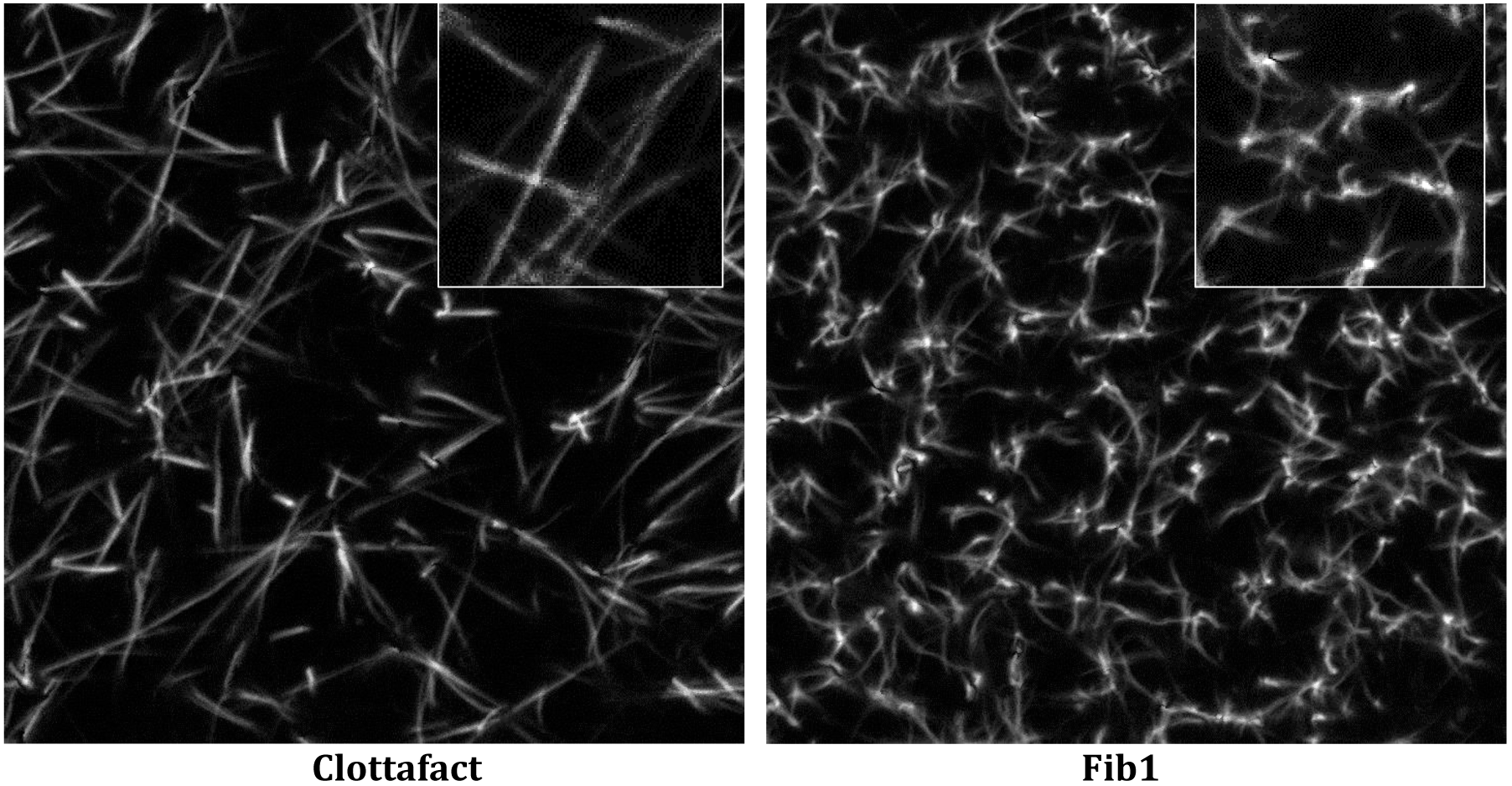
Qualitative geometry of the fibrin networks. Smooth Manifolds were extracted as described in material and methods from confocal microscopy stacks. Clots were formed using 3 mg/ml and coagulated using 0.1 IU/ml of thrombin. Frames are 67.5 μm wide. The insets are x2 enlargements of a portion of the original SME image (12×12 μm).

Another major difference between those two networks is highlighted in the insets of figure 5. Clottafact’s fibrin shows a rather small number of branches per branching node, typically looking like the contact point between needle-like fibers while Fib1 shows many star-like patterns, with many asymmetrical branches per node expanding in thin fibrin threads.

## 4 Discussion

The first aim of this article is to investigate whether or not fibrinogen dispersity influences the ensuing fibrin ultrastructure and how important it is with respect to compositionally-induced changes. A possibility to obtain such products would have been to replicate the fractionation process proposed by Huang et al (11), followed by a complete determination of the fractionated fibrinogen composition. Unfortunately, this fibrinogen fractionation method has been shown to produce important amounts of in-column degradation products (13) which contaminated the elution peaks of the undegraded material. In consequence, we chose to test existing commercial fibrinogens preparations for dispersity. Among those we tested, only Clottafact showed a close-to-monodisperse profile. For the second, polydisperse fibrinogen, we chose Fib1 fibrinogen which is the *de facto* reference since it has been used in the large majority of works dealing with purified fibrin structure and rheology (see SI1).

In the first part of this work we determined the Fib1 and Clottafact compositions by measuring their FXIII, fibronectin, α,β and γ intact-chains content, γ/γ’ ratio as well as their N-glycosylation and post-translational modifications (see table 1). To evaluate if the variations in composition observed between Fib1 and Clottafact may have a significant impact on fibrin structure, we compiled the compositional effects on fibrin structure observed in the literature. This compilation (table S3.1 in SI3) shows that the measured Clottafact-Fib1 compositional differences are (very) small compared to the compositional range of variation necessary to obtain measurable structural changes in fibrin structure. So, the slight compositional variance between Clottafact and Fib1 cannot explain the large structural differences observed between Clottafact and Fib1 fibrins. Those structural differences can then only be the consequence of the significant quantity (4% in mass) of large aggregates found in Fib1 since Clottafact does not possess such aggregates. Surprisingly, aggregates-induced structural differences are larger than those induced by compositional changes under constant reaction conditions, as shown in SI3 table SI3.1. Therefore, the presence of aggregates is a major determinant of fibrin polymerization and structure and its effect persists at all concentrations used in this work, from 0.5 mg/mL to 3 mg/mL.

To explain this phenomenon, we start from fig. 5 which shows that a central difference between Clottafact and Fib1 microstructure is the number of fibers/branches emerging from a branching node. A branching node in Fib1 shows typically 6 or more branches while a branching node in Clottafact rarely exceeds 4 branches. Branching in Clottafact looks like fortuitous contacts and adhesion between already formed needle-like fibers in a very similar way to the Mikado model of fibrin networks proposed by Lang et al. (28). Conversely in Fib1, each branching looks like an aggregate and seems to act as a nucleus from which several fibres grow in a rather random fashion. To confirm that aggregates may act as nucleating centres, the average distance *d* between them (which is also the pore size) can be estimated and compared with the *measured* pore sizes. Aggregates are the centre of spheres of radius *d*/2: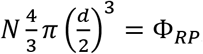, where *N* is the number concentration and Φ_RP_ is the random packing fraction for monodisperse hard spheres. So, 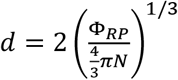, which gives a pore size of 4μm for 4% in 4 mass of aggregates in 3g/L of Fib1, value in remarkable agreement (for such a simple model) with the 3μm experimentally measured pore size (see SI7 fig. SI10Left).

Because most Fib1 fibers appear to nucleate from those aggregates, the larger the aggregates content is, the more branches and hence the more fibres. Since the total fibrinogen mass is conserved, it further derives that the larger the aggregates content is, the smaller the number of protofibrils should be in the ensuing fibrin. Since the Clottafact and Fib1 have practically identical compositions, this hypothesis can be checked by mixing increasing quantities of Fib1 with Clottafact, experiment which amounts to increasing the aggregates content. Fig. 6A shows a large (~100%) linear decrease in the number of protofibrils per fiber as a function of the aggregate (~Fib1) content, as expected from the simple argument presented above. Confocal images SI8 fig. S11 of the same mixtures show also a progressive change from straight fibers to rather curved fibers linking bright aggregates, supporting the above explanation.

**Figure 6 A:**
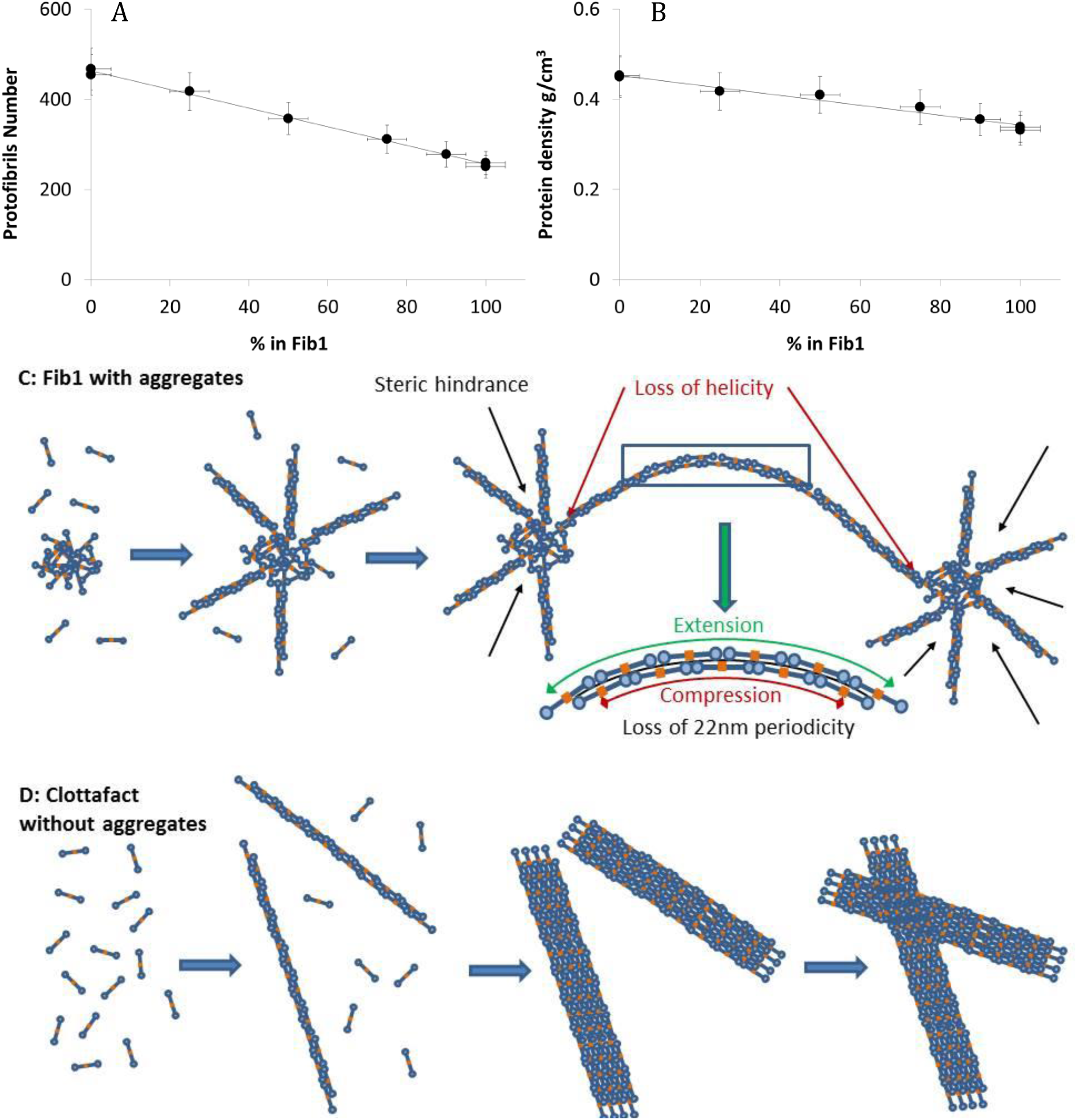
Number of protofibrils per fiber vs content in Fib1. B: Fibrin Fibers internal density as a function of Fib1 content. The content in Fib1 is proportional to the amount of aggregates in solution. C: proposed fibrin formation mechanism in the presence of aggregates (Fib1). D: Proposed fibrin formation mechanism in the absence of aggregates (Clottafact).

Fig 6B shows that the internal density of the fibers is also significantly modified by the presence of aggregates, but in a less dramatic fashion than the number of protofibrils. As deduced from the confocal images above, a plausible hypothesis is that a large part of the Fib1 fibers actually grow from activated fibrinogen present in the aggregates, hindering the lateral growth through several phenomena. First, there will be steric hindrance because of the local presence of the aggregate and of the several neighboring fibers as shown in fig 6C. Second, to link between aggregates, the fibers can stick upon each other or the fiber can bend. In the latter case, such a bending will discourage lateral growth because the 22.5 nm periodicity will be partly lost. Finally, recent results suggest that it is the twisted conformation of both fibrinogen monomers and protofibrils that promotes the assembly of protofibrils into fibres, in agreement with earlier observations by Weisel’s group (29). Therefore, aggregates could disrupt the initial fibres natural helical geometry (12), forbidding a proper local lateral aggregation, generating defects or holes in the structure of the fibres. All of these possible mechanisms lead to the same result: a net decrease of the internal density of the fibres. Conversely, in the absence of aggregates, ideal needle-like fibres can grow and then stick together when they meet. So, once again, the larger the aggregates content is, the larger the number of perturbed, low density fibres will be, giving an average decrease in fibre density as observed in fig. 6B above. This interpretation also explains the absence of well-defined lateral structure in Fib1 fibers as observed in the SAXS experiments.

### Conclusions

We have first shown that Clottafact’s fibrinogen is composed almost exclusively of fibrinogen monomers whereas Fib1 possess a significant amount of large aggregates. Conversely, the detailed physico-chemical analyses of Clottafact and Fib1 show that their compositions are sufficiently similar not to influence the structure of the ensuing fibrins. Small angle X-ray scattering demonstrates that the internal structure of Clottafact’s fibrin fibers is close to crystalline both laterally and longitudinally with a longitudinal persistence length of several microns. Conversely, SAXS shows that Fib1 fibers are much more disorganized with no perceptible lateral organization and a longitudinal persistence length of a few hundreds of nanometers. Those nano-structural observations are confirmed at the microscale by direct confocal imaging. Clottafact fibrin displays long, needle-like fibers while Fib1 fibrin shows short and curved fibers in agreement with the SAXS persistence length measurements. Those results are further strengthened by spectrophotometry measurements which confirm the much larger density of the Clottafact fibers with respect to Fib1 ones. Those structural differences hold at all studied concentrations including physiological ones and demonstrate that the size dispersity of fibrinogen in purified systems is one of the main parameters determining fibrin fibers ultrastructure, parameter that has been mostly overlooked. Indeed, the effect of modifying the dispersity is surprisingly large, larger than changing any other aspect of the fibrinogen preparation.

Those results open outstanding questions regarding the precise mechanisms by which the presence of aggregates can influence to such an extent the polymerization of fibrin and, therefore, concerning fibrin polymerization itself. Furthermore, given the extensive structural modifications accompanying the presence of aggregates, the mechanical properties of fibrin clots formed from monodisperse fibrinogens may be very significantly different from the polydisperse ones and could be of paramount interest in tissue engineering on fibrin scaffolds as well as for the design of fibrin glues.

Finally, the actual amount of aggregates in circulating fibrinogen is not known as only a few studies investigated the multiscale structure of fibrin formed in plasma. Such investigations could be of outstanding interest in hemostasis, in particular in relation to thrombosis related diseases.

## Authors’ contributions

XG, LS and FC conducted and analyzed the experiments in small angle X-ray scattering, multi-wavelength spectrophotometry, dynamic light scattering, confocal microscopy and gel filtration chromatography. GC and NB conducted and analyzed the experiments in reverse phase chromatography, mass spectrometry and ELISA dosages. FC, BP and ZT conceived and directed the research. XG, BP and FC wrote the manuscript. All authors contributed to the final version of the manuscript.

## Acknowledgment

The authors gratefully acknowledge the help of Marie-Claire Dagher and Caroline Mas for gel-filtration experiments, Christophe Travelet for DLS experiments, Denis Roux, Emmanuelle Bigo and Michael Sztucki (local contact) for the SAXS experiments performed on beamline ID02 (ESRF, Grenoble, France). The SEC-MALLS-RI was performed at the Grenoble Instruct-ERIC Center within the Grenoble Partnership for Structural Biology. The authors are also grateful to Marguerite Rinaudo and Marie-Hélène André for a careful reading of the manuscript.

Xabel García was funded by the LabEx Tec 21 (Investissements d’Avenir - grant agreement n°ANR-11-LABX-0030).

